# G-Dash: A Genome Dashboard Integrating Modeling and Informatics

**DOI:** 10.1101/501874

**Authors:** Zilong Li, Ran Sun, Thomas C. Bishop

**Affiliations:** Louisiana Tech University, Computational Analysis and modeling; Louisiana Tech University, Engineering Physics; Louisiana Tech University, Department of Chemistry and Physics

**Keywords:** Genome Browser, genome dashboard, gene editing, next generation sequencing, genetic engineering, chromatin structure, chromatin modeling

## Abstract

Genomics is a sequence based informatics science and a structure based molecular material science. There are few tools available that unite these approaches in a scientifically robust manner. Here we describe G-Dash, a web based prototype of a genomics dashboard, specifically designed to integrate informatics and 3D material studies of chromatin. G-Dash unites our Interactive Chromatin Modeling(ICM) tools with the Biodalliance genome browser and the JSMol molecular viewer to rapidly fold any DNA sequence into atomic or coarse-grained models of DNA, nucleosomes or chromatin. As a chromatin modeling tool, G-Dash enables users to specify nucleosome positions from various experimental or theoretical sources, interactively manipulate nucleosomes, and assign different conformational states to each nucleosome. As an informatics tool, data associated with 3D structures are displayed as tracks in a genome browser. The exchange of data between informatics and structure is bi-directional so any informatics track can inform a molecular structure (e.g. color by function) and structure features can be displayed as informatics tracks in a genome browser(e.g. Roll, Slide, or Twist). As a sample application, models of the CHA1 promoter based on experimentally determined nucleosome positions are explored with G-Dash. Steric clashes and DNA knotting are observed but can be resolved with G-Dash’s minimal coarse-grained model without significant variation in structure. Results raise questions about the interpretation of nucleosome positioning data and promoter structures. In this regard, G-Dash is a novel tool for investigating structure-function relationships for regions of the genome ranging from base pairs to chromosomes and for generating, validating and testing mechanistic hypotheses. http://dna.engr.latech.edu/∼gdash/GDash-landing-page

## INTRODUCTION

Chromatin is the biomaterial that contains the genome in all higher organisms. Genomics is fundamentally a DNA sequence based information science and a 3D based material science. Determination of chromatin structure is challenging for both experimentalists and theorists[1]. There is no consensus for the structure of chromatin[2], but there is a wealth of structure, informatics, and functional data. 3D structure based experiments provide atomic or near atomic data for nucleosome and nucleosome arrays[3-9]. Cross-linking and sequencing based methods yield the gross structure of entire genomes[10-17]. Global efforts such as The 1000 Genomes[18; 19], ENCODE[20-22] and the 4D Nuclesome[23] projects provide reference standards for informatics analysis. Coupled with next generation sequencing(NGS) and genome wide association(GWA) studies these reference standards enable individual labs to link chromatin reprogramming with disease and altered gene expression, e.g.[24-27]. However, a significant challenge in chromatin structural biology is unifying this data to develop and validate structure-function relationships or hypotheses genomic mechanisms of action.

There exists a growing collection of tools for low resolution chromatin modeling based on Hi-C analysis[15; 28], but there are few if any tools that directly link informatics data with atomic or coarse-grained modeling. Researchers often rely on conceptual diagrams to describe genomic mechanisms. These diagrams may fail to account for known structural details. Likewise, even though atomic and coarse-grained models of chromatin are rapidly maturing[29; 30], modelers are faced with the challenge of identifying relevant experimental and biological data during study design and interpretation phases. Modelers may fail to account for important biologic information.

The goal of G-Dash is to promote the convergence of informatics and structural studies of chromatin through the development of genome dashboards. A dashboard is a console that manages information and provides controllers for navigating the physical world. A “genome dashboard” merges bioinformatics and structural studies of genomes. Merging data from different sources (data unification) is based on the idea that DNA is the common thread in chromatin structural biology and is achieved through laboratory (Cartesian coordinate) and material (internal coordinate) reference frame representations of DNA as a space curve.

Here we present G-Dash as a prototype genome dashboard for modeling chromatin. Below we describe the mathematical theory and code design principals that are the foundation of the genome dashboard concept. We then introduce G-Dash and demonstrate a simple usage scenario using the CHA1 promoter as an example. We conclude with a summary of G-Dash capabilities and outline future possibilities.

## METHODS

### Theory

#### Geometry

Chromatin is a protein-DNA complex whose structure is synonymous with the 3D space curve of DNA contained within it. In a laboratory reference frame, a continuous space curve has centerline,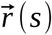, and director frames embedded in the underlying material, represented by matrix 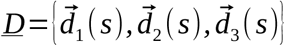 [31]. For DNA, a natural choice is to align the three directors with the major and minor grooves and the axis of DNA. An equivalent description of the curve is based on the director frames themselves. This description is a material reference frame description that captures the translations and rotations connecting one director frame to the next, represented here by 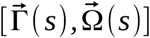. The two representations are related by the expressions 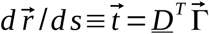 and 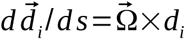, where 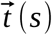 is the unnormalized tangent, *<u>D</u>*^*T*^ is the transpose of the matrix whose columns are the directors, and 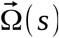 is the vector representing the instantaneous axis of rotation of the director frame located along the curve at position, s. The Frenet-Serret Tangent, Normal, Binormal (TNB) description can be obtained from just a centerline[32]. For a TNB description of DNA assumptions about the distribution of twist (for example uniform twist) and shear (for example shear free) have to be employed to construct director frames that align with base pairs. In this manner the TNB approach allows a 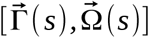 representation to be obtained even when director frame data,*<u>D</u>*(*s*), is missing.

The mathematical foundation of the genome dashboard concept is that 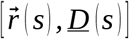 and 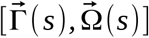 are equivalent descriptions of the conformation of a space curve, denoted simply as *C* (*s*). Converting between representations requires either an integration or a differentiation. The two representations provide a basis for merging structure and informatics data. Data unification is thus achieved by indexing all structure data *C* (*s*) and informatics tracks *T* (*s*) by chromosome coordinate *s*.

##### Free DNA

DNA conformation C(s) is at best a base pair discrete approximation to a continuous space curve. (See pioneering efforts by Callidine and Drew[33] and recent efforts in [34-37].) The base pair step parameters[38; 39] and associated algorithms[40; 41] provide established conventions for describing double and single stranded DNA as a space curve at atomic resolution. A sequence specific di-nucleotide accurate model of dsDNA in the 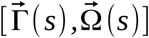 (helical parameter) representation can be obtained from x-ray[42; 43] or molecular dynamics[29; 44] studies. The 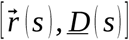 description is obtained by integrating 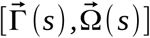 as noted above. Thermal motion can be introduced as random variations of the helical parameters scaled according to measured values of standard deviation[45]. There are two widely used tools for DNA helical parameter analysis. 3DNA[46] uses an Euler angle method and a mid-step plane approximation[40]. Curves+[47] uses a Cayley parameter based method[37]. These and other tools[48; 49] are extremely fast. Helical parameter values obtained from the two methods are known to differ[50] Differences may arise from two sources. The assignment of director frames to base pairs may differ. However, the methods for assigning director frames are well defined[51], so these differences should only occur for significant deviations from ideal pairing. The other source of differences is method dependent. Helical parameter values obtained from the Euler based method, with its mid-step plane approximation, and from the Cayley parameter method differ even when the director frames used for the calculations are identical. Thus, values obtained from one method should not be interchanged with the other.

##### Masked DNA

Histones or other external agents can alter the material properties of a contiguous length of DNA from *s* to *s*+*N*. We label this a Mask, *M* (*s*). Any number of identical or unique Masks may be associated with a sequence of DNA. In the material reference frame the conformation of the masked DNA is denoted as 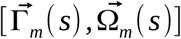 and spans *N* base pairs. As the name suggests, the Mask simply replaces the 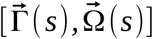 values associated with free DNA. In terms of Masks, a nucleosome is a DNA superhelix *and* histones. The histones can be represented as a single entity (sphere, cylinder, ellipsoid), a collection of beads, or an all-atom model. Docking the histones to the superhelix is achieved with the same methods and tools used for describing the relative rotations and translations of base pairs. A Mask thus has two components: altered material properties compared to free DNA (conformation and/or flexibility) and docking parameters. If the Mask is a rigid entity both components are described by fixed lists of internal and Cartesian coordinates. The Cartesian coordinate representation of C(s) requires only a single translation and rotation to position each rigidly Masked element in the laboratory reference frame. Linker DNA connecting Masks is constructed as free DNA.

##### Chromatin Folding

We define chromatin folding as the physical process by which proteins or other external agents associate with DNA and alter its material properties (conformation, dynamics, flexibility, or chemical properties). DNA methylation is considered a Mask by this definition since it does not change DNA sequence. In the context of a genome dashboard, chromatin folding is an informatics problem of describing all the unique Masks and tracking their locations along a sequence of DNA. G-Dash demonstrates that such an inventory of Masks can be maintained and converted to 3D structures from single base pairs or entire chromosomes in real time. In this manner, genome dashboards enable users to both define and navigate chromatin folding energy landscapes.

#### Energy

There is no way of knowing if a model assembled in the material representation, 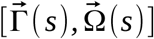, will avoid steric clash or knotting in the laboratory representation,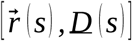. We utilize LAMMPS to relax G-Dash structures using an approximate coarse-grained model. The purpose of this model is only to solve steric clashes and knotting as rapidly as possible in LAMMPS without causing significant variation from the initial structure. It is not intended for thermodynamic sampling. The model parameters yield a chromatin structure that is compatible with[52] [53]. The current model in G-Dash does not include histone tails and includes only three types of beads: Free DNA, Nucleosome DNA, and Octasomes, as shown in Figure 1. We utilize harmonic bond functions *E*= *K* (*r* − *r*_0_)^2^ and define three bond types to describe: bonds between 1) Free DNA, 2) Octasomes and Nucleosomal DNA, and 3) Nucleosome DNA beads. The second and third bond types maintain nucleosome comformation during relaxation. A harmonic angle term captures DNA stiffness *E*= *K* (Θ−Θ_0_)^2^ based on the angle between adjacent DNA-DNA bonds. Finally, a soft pair repulsion, 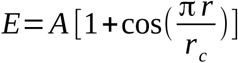 is used to push apart overlapping atoms. No effort is made to capture electrostatic or van der Waals interactions in this model. The parameter values utilized are summarized in Table 1.

**Table 1:**
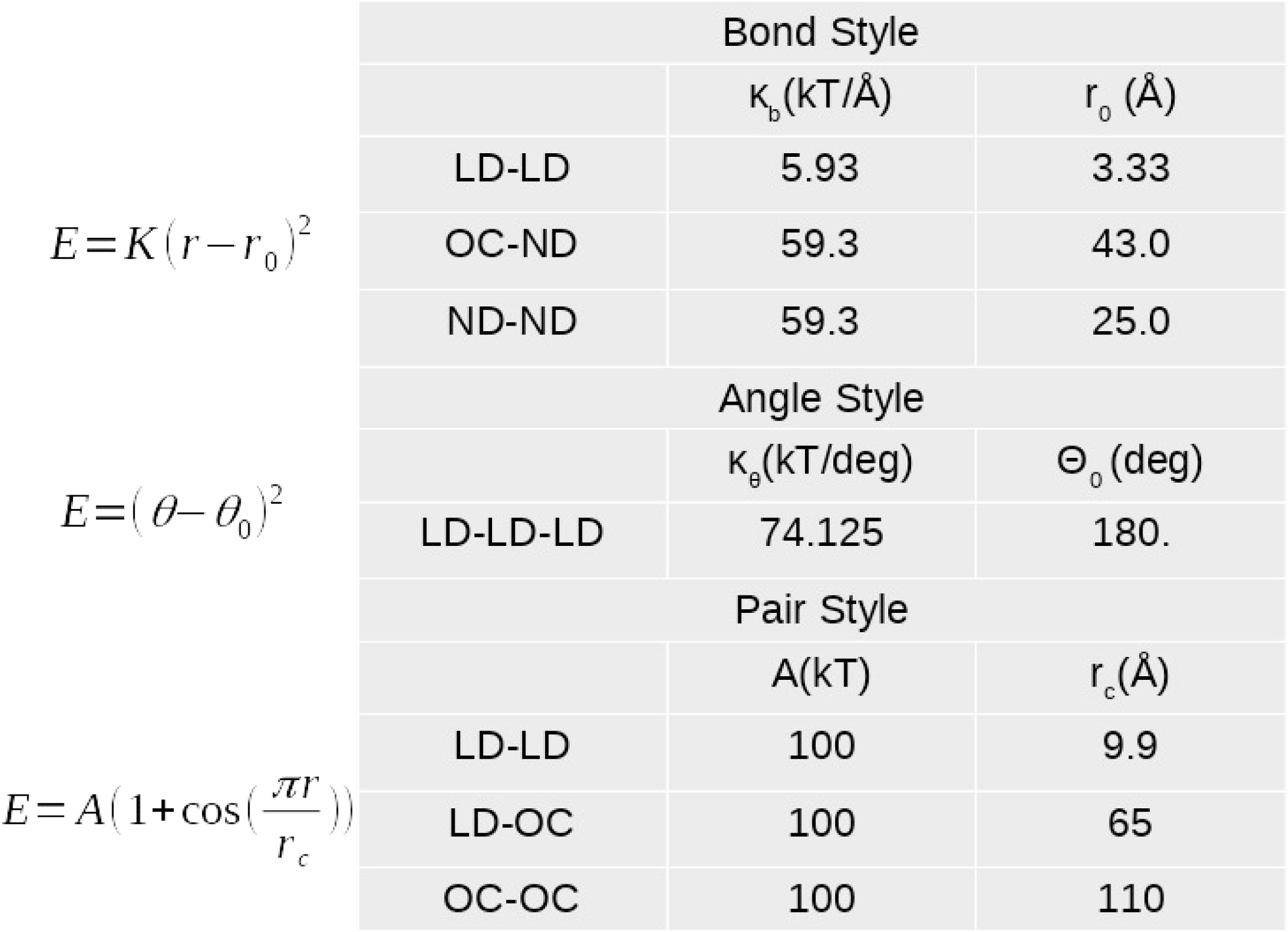
Energy Terms for the Coarse-Grained Model. The coarse-grained model uses harmonic bond and angle functions and a soft-core repulsion term available in LAMMPS. Parameter values for each energy function and atom type are as listed. These values are chosen to preserve structure while resolving steric clash and knotting rather than for thermodynamic accuracy.

**Figure 1:**
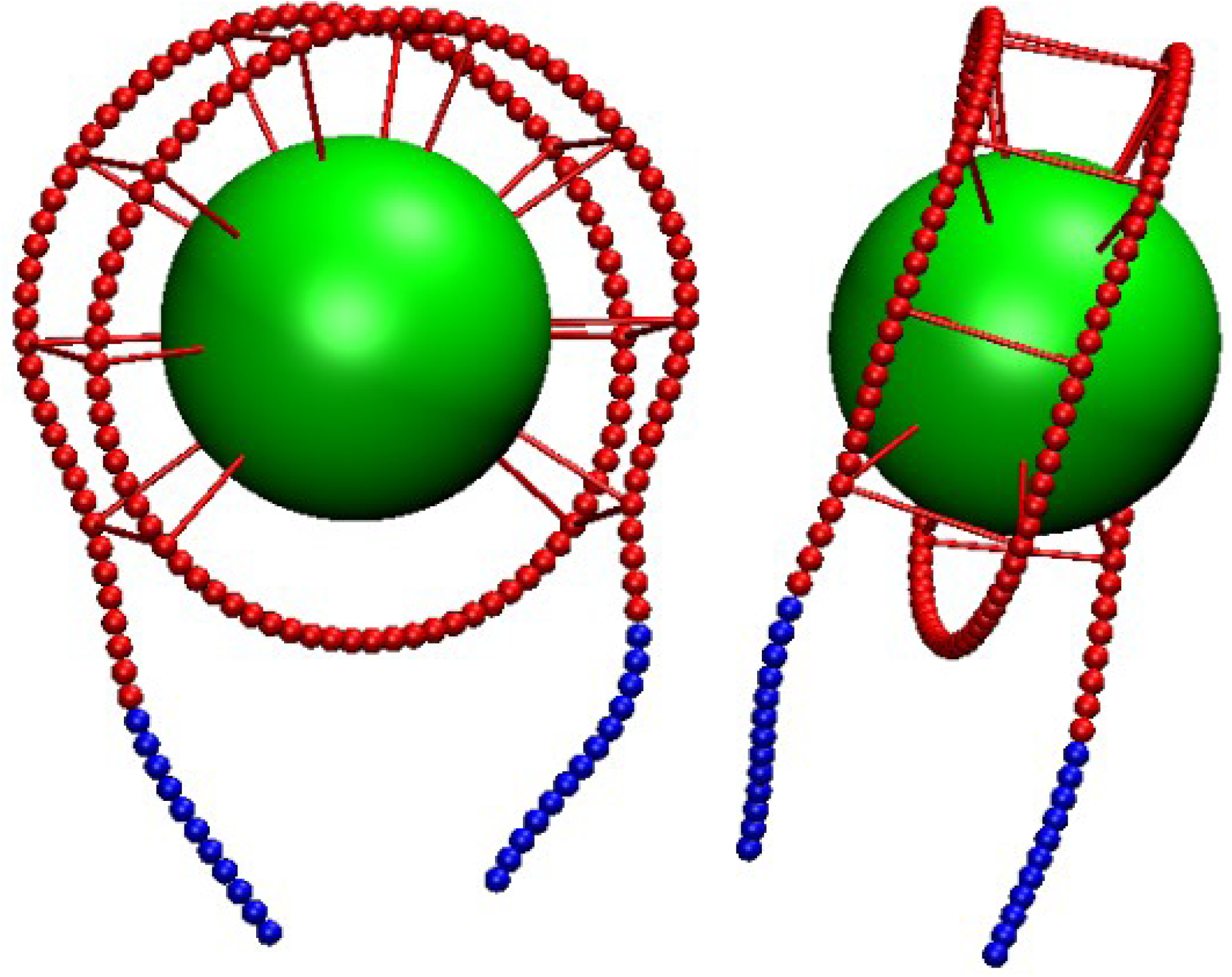
Coarse-Grained Nuclesome Model: The nucleosome model used during structure relaxation is a minimalistic model designed to preserve nucleosome structure. It consists of three “atom” types: (1) free DNA (blue beads), (2) nucleosomal DNA (red beads), and (3) the histone octamer (green bead). There are three “bond” types: (1)-(1), (2)-(3) and (2)-(2). The latter fixes the pitch of the DNA superhelix with bonds between adjacent atoms of the gyres.

### Code Design

Genome dashboards can be efficiently designed using Model-View-Controller(MVC) design principals that enforce separation of Concerns(SoC)[54], Figure 2. This ensures that the dashboard’s functionalities overlap as little as possible, are replaceable and extensible. The MVC idea separates the Model, the Views, and the Controller, thus allowing independent development of each.

**Figure 2:**
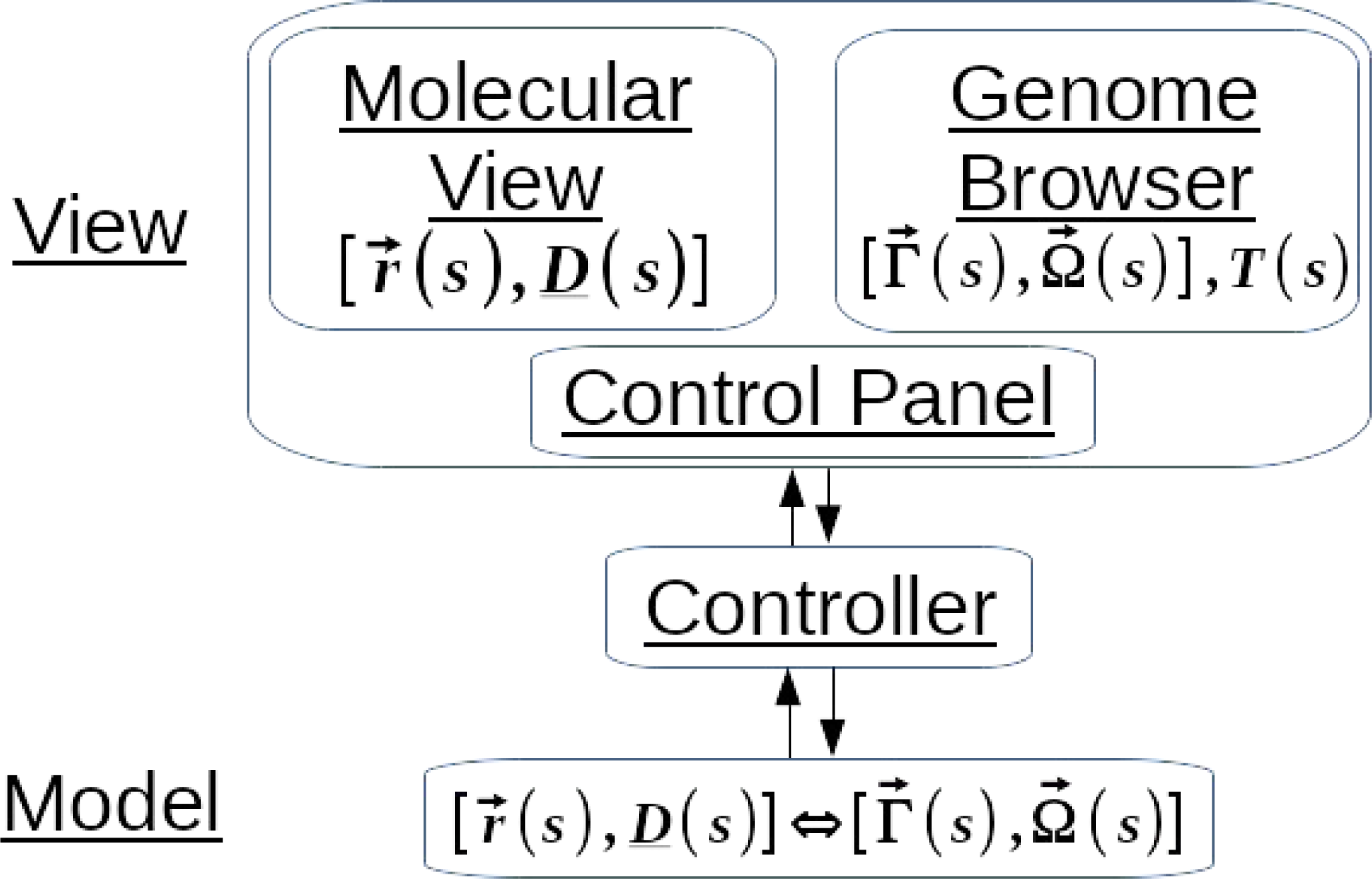
Model-View-Controller(MVC) Design: A genome dashboard is a finite state machine that can be efficiently developed with a Model-View-Controller(MVC) design philosophy. The Model includes: Cartesian coordinate and material reference frame descriptions of DNA as the common thread, an inventory of Masks, and the necessary logic and routing to account for the Mask when coverting between the representations. The Views include: a Genome Browser, a Molecular Viewer and and a Control Panel. G-Dash uses Biodalliance and JSmol for the browser and viewer elements, respectively. The Control Panel is specific to G-Dash. And appears as extension of Biodalliance, the Controller manages the exchange of data between Model and Views.

#### Model

Model in the MVC schema is the data and related logic. For a genome dashboard, the Model includes the 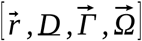 representations of DNA as a base pair discrete space curve overlaid with Masks, *M* (*s*), the associated track data, *T* (*s*) and tools for converting between representations. In general, the rotations and translations associated with a Mask may be large. G-Dash currently uses the Euler-based algorithm for conversions between 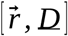 and 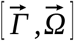, which assumes a mid-step plane approximation. This method is restricted to small bend values suitable for DNA[40]. A Cayley parameter based method is a more appropriate choice since this method is not restricted by these approximations. If the Masks are all rigid entities they can be leveraged to improve performance. For example, representing a nucleosome as a Mask with one bead for both DNA and histones and a single large deformation of the path of DNA reduces computational and data costs by approximately *n*\*146, where *n* is the number of nucleosomes containing 147 base pairs. G-Dash currently maintains a base pair discrete model throughout that does not exploit these performance enhancements.

#### View

View in the MVC schema includes the Control Panel (CP), a 3D Molecular Viewer(MV) and a Genome Browser(GB).

For the Molecular Viewer, G-Dash utilizes JSmol. In G-Dash, all-atom models are generated with 3DNA and parameterized with Amber’s tleap modules. The models include parmtop, crd and pdb formatted files that can be downloaded by the user to initiate modeling on their own computing resources. JSmol displays the pdb file using JSmol’s cartoon style by default with adenines(A) colored red, thymines(T) blue, guanines(G) green and cytosines(C) yellow. Coarse-grained models are stored in an xyz formatted file. The DNA space curve has Center Atoms, CA, director frame end points as H1, H2, H3 atoms, and octasome cores as OC atoms. We have applied the following rules for representing coarse-grained models in JSmol. DNA is represented by small beads and nucleosomes by large beads. A small green bead represents the start of the structure, and yellow beads represent intermediate base pairs. Large red beads indicate an unresolved steric clash in models containing less than 30,000 base pairs. For larger models and nucleosomes lacking steric clash, the nucleosomes are represented by blue beads. Steric clash is not monitored in large structures to maintain the interactive nature of G-Dash. For all models JSmol is fully functional so users can save files, change colors or representation schemes etc…

For the Genome Browser, G-Dash utilizes the Biodalliance Genome Browser[55]. As embedded in G-Dash, Biodalliance allows users to choose the SerCer3 assembly of the yeast genome or the hg19 and hg38 assembles of the human genome. G-Dash can be configured to use other public or private genome assemblies. For each genome, users are able to enter the chromosome coordinates of interest or a search term to jump to a desired location. For example, entering CHA1 for the sacCer3 assembly jumps to chromosome coordinate “III:5,798..26,880”. In a genome browser all data are displayed as tracks. In G-Dash, several default tracks are pre-selected, but users can add tracks from either public or private resources[56]. For sacCer3 the default tracks include “Occ”, “Genes”, “Count” and “Peak” and represent data obtained from [57].

Biodalliance supports bigwig, 2bit and other common data formats. Users can manipulate the style, color, and max/min values associated with any track. In G-Dash, users select a specific DNA sequence for modeling using the sequence selector (a yellow bar) in the top track of the genome browser. G-Dash automatically updates structural-informatics tracks in the genome browser whenever a model is updated. Structural-informatics tracks are a unique feature of the genome dashboard concept. These tracks represent features extracted from the 3D model. In G-Dash structural-informatics tracks currently include “Energy”,”Nucleosomes”,“Shift”, “Slide”, “Rise”, “Tilt”, “Roll”, and “Twist”, tracks.

For the Control Panel, a graphical user interface has been appended to the Genome Browser. The Control Panel contains the Nucleosome Energy Landscape introduced in ICM[45]. The Nucleosome Energy Landscape is a specific instance of a more general Mask Widget that both represents and enables manipulation of Masks. In the stable version of G-Dash users can manipulate nucleosome positions and types. Additional controls affect predetermined computations such as buttons are described in more detail in the Usage Scenarios section.

#### Controller

Controller in the MVC scheme is the interface between View and Model that processes all user input. G-Dash relies on multiple data and language standards to achieve bi-directional exchange of data between informatics and structure so that any informatics track can inform a molecular structure and structural features can be extracted as informatics tracks. G-Dash uses HTML as the front end, JavaScript and JSON(JavaScript Object Notation) for HTML functions and for embedding Biodalliance, and PHP to pass data from web pages to modeling units that are coded as unix and vmd scripts, FORTRAN, Python, and C. G-Dash utilizes bigwig[58] and 2bit genomics formats to exchange track data with Biodalliance. A Python based implementation of the Model as an integrated compute kernel is under development.

## USAGE SCENARIOS

A genome dashboard, like any dashboard, enables a user to navigate a physical world that appears in one or more windows. The navigator may choose to utilize informatics data, observations of the physical world, autopilot or any combination of these to achieve a desired outcome. An experienced pilot maintains situational awareness and knows which data sources should be monitored or ignored to achieve a desired outcome. Below we describe some simple usage scenarios, and the outcomes that can be achieved with G-Dash.

Additional details including video demonstration included as help on G-Dash landingpage.

### From Informatics to 3D Model

One usage modality of G-Dash is the generation of a 3D model based on informatics data. For this usage modality, tracks *T* (*s*), and Masks *M* (*s*) describing selected nucleosome models[59; 60], together with 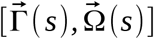 data for free DNA obtained from [29] are used to generate 3D models. Below we describe how to make all-atom or coarse-grained models of DNA, mono-nucleosmes or chromatin with this usage modality.

### Free DNA Modeling

As with ICM, all-atom or coarse-grained models of DNA required only a sequence and temperature to be specified. G-Dash reads DNA sequence information directly from Biodalliance. Users select the sequence using the sequence selector(yellow bar in genome browser window), and the chromosome coordinates of the selected sequence appear in the Control Panel as the Model Coordinates. Temperature (“T” in the Control Panel), in degrees Kelvin, determines the amount of thermal variation to be added to the helical parameters for the regions of free DNA. G-Dash has four options, 0, 100, 200, and 300, available as a drop down menu in the Control Panel. The default value is zero which means sequence specific average values of 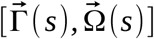 as observed in [29] are used. The Temperature function works for both free DNA and chromatin. However only the DNA that is not Masked is subject to thermal variations. As we reported previously [59] random fluctuations added to the nucleosomal DNA helical parameters are sufficient to destroy the nucleosome superhelix[59]. For this reason thermal variations are not added to any region of DNA that is masked. The Free DNA button in G-Dash will generate a single 3D structure or ensemble containing ten different sequence specific thermal variants. In the development version of G-Dash, an all-atom model can be generated using the “All Atom” button in the Control Panel for any single coarse-grained models.

### Nucleosome Modeling

The stable version of G-Dash includes Masks that represent various conformations of the nucleosome superfamily of states[60]. All-atom and coarse-grained modeling is allowed for nucleosomes and is controlled by the Nucleosome Widget in G-Dash. A general purpose genome dashboard should include a Mask Widget. The idea is the same as the Nucleosome Widget described below.

#### Nucleosome Widget

The Nucleosome Widget represents and controls the location and type of nucleosomes in a Nucleosome Energy Landscape, see Figure 3. The Nucleosome Energy Landscape was described in ICM[45] and is used to postion nucleosomes. In G-Dash users may position nucleosomes based on any informatics track. Thus, the Nucleosome Energy Landscape may serve as only a reference for manipulating nucleosome positions and types. G-Dash users are able to add, delete, move or alter nucleosome types using the Nucleosome Widget by clicking on the corresponding block in the Nucleosome Energy Landscape. The block turns green when selected and can be used to identify a nucleosome in a 3D model. Bi-directional data exchange was not supported in ICM-web so these functionalities were not available.

**Figure 3:**
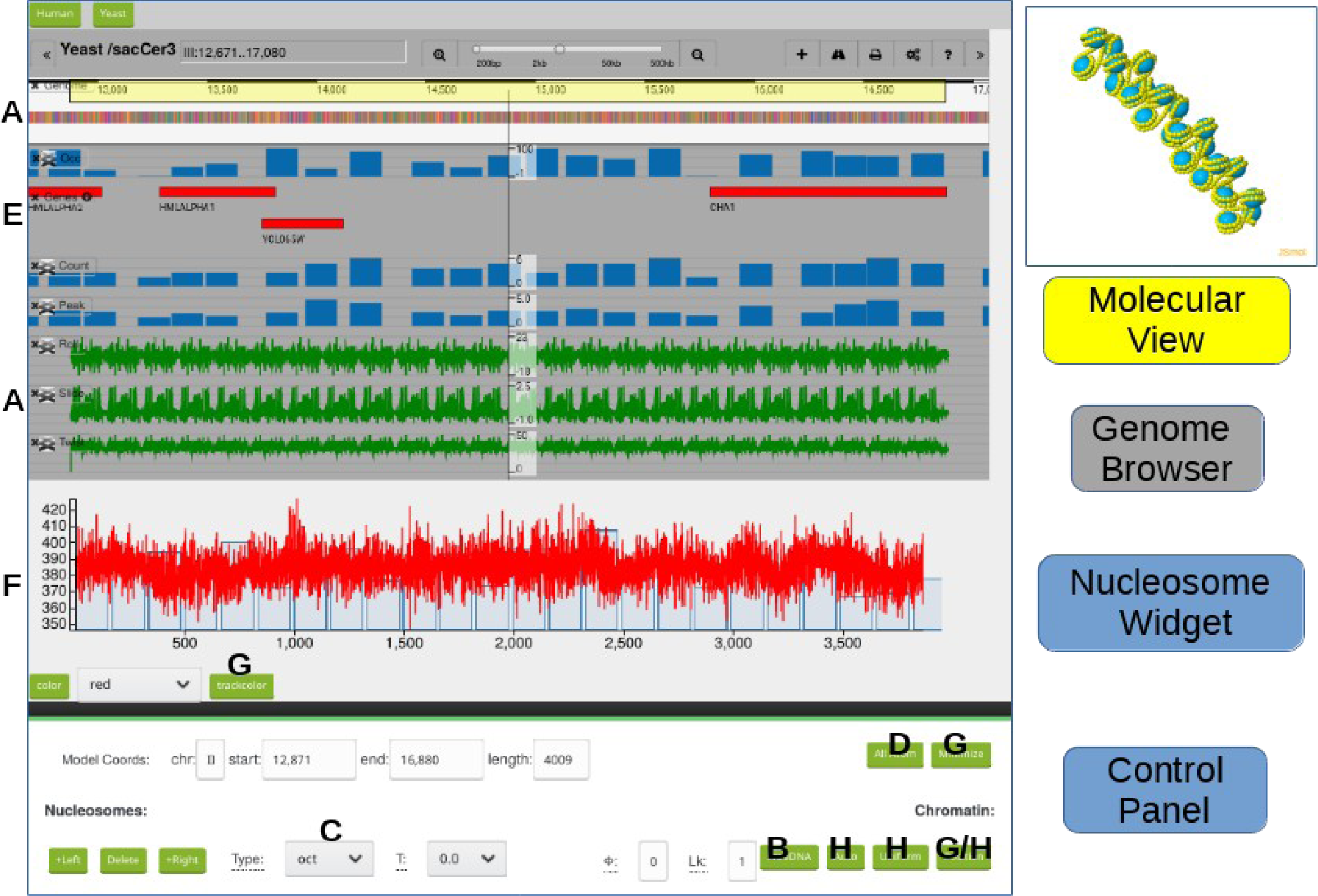
Summary of G-Dash Views: G-Dash contains an embedded genome browser (Biodalliance), a Nucleosome Widget, a Control Panel, and a Molecular Viewer (JSmol). The Nucleosome Widget is a component of the Control Panel that displays a Nucleosome Energy Landscape and allows nucleosomes to be individually manipulated. See text for additional descriptions of Control Panel elements. Letters corresponds to descriptions in text and items in Figure 4.

#### All-atom Nucleosome models

G-Dash will make an all-atom mono-nucleosome model for any nucleosome selected from the Nucleosome Energy Landscape by clicking the “All Atom” button in the Nucleosome Widget, Figure 4 D. The models are based on the 1KX5 x-ray structure[61] and include Amber formatted parmtop, crd, and pdb files that can be downloaded for computational studies by the user. For these models a DNA superhelix is constructured for the selected sequence of DNA and docked onto 1KX5’s histone octamer. Coupled with high performance high throughput workflows[62] and our iBIOMES-Lite[63] database of nucleosome simulations[64], a software ecosystem now exists for overnight comparative molecular dynamics simulations of nucleosomes.

**Figure 4:**
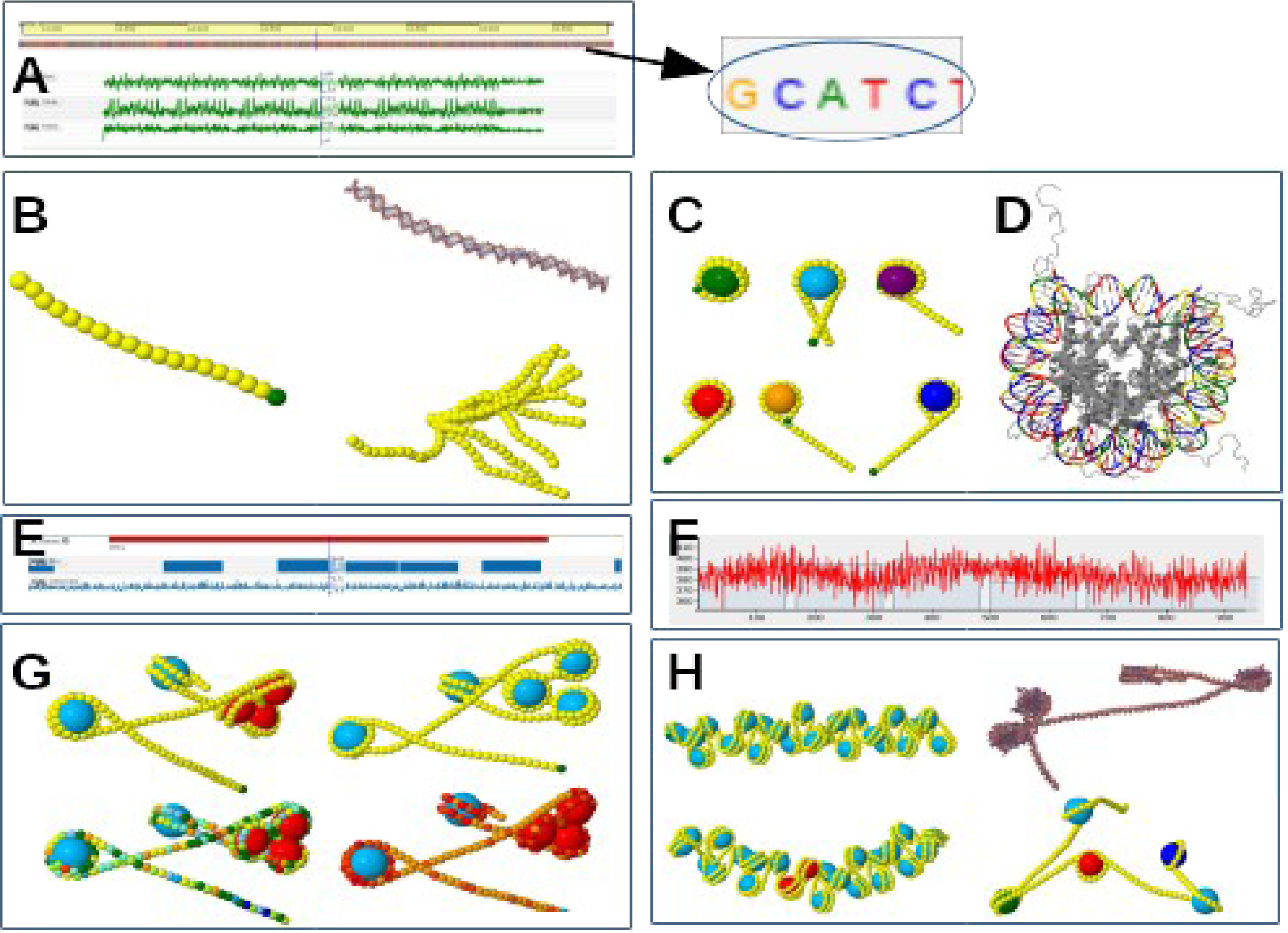
Typical Usages of G-Dash. A) A sequence is selected for modeling with the yellow highlighter in the sequence track of the genome browser (yellow bar). B) A single sequence specific coarse-grained or all-atom model of DNA or ensemble of conformations representing thermal variations can be generated and displayed. C) Different conformations of the nucleosme superfamily of states can be assigned to specific locations using informatics tracks E) or the Nucleosome Widget F). D) An all-atom model for any single nucleosome can also be generated. G) The default coloring of coarse-grained nucleosome models uses small yellow beads for DNA and large blue beads for the histones. Steric clash is indicated by red nucleosmes. Informatics data can be mapped on the coarse-grained models. H) Options for generating specific distributions of nucleosomes along the DNA are provided. The development version supports conversion of coarse-grained models to all-atom models.

#### Nucleosome Superfamily

Besides the standard nucleosome(octasome), sub-octasome and super-octasome states of the nucleosome also exist. G-Dash gives the option of utilizing different nucleosome states in a single coarse-grained model of chromatin. The stable version of G-Dash supports manual positioning of octasomes(oct), hexasomes(hex1,hex2), tetrasomes(tet), and chromatosomes(chrmat). An informatics track can be loaded into a genome dashboard to track these structural variations. These features were not possible with ICM-Web.

### Chromatin Modeling

Given the ability to model free DNA and nucleosomes, chromatin modeling is achieved by assembling these building blocks. Chromatin is linker (free) DNA interspersed with nucleosome Masks *M* (*s*). To locate the nucleosome Masks, users can utilize any single informatics track or use G-Dash’s positioning tools as described below. Whenever a model is created structural-informatics tracks in the genome browser are automatically updated. Figure 4, H. Collectively these tools provide a novel means of investigating structure-function relationships.

#### Automatic positions

The auto option automatically places nucleosomes in the energy landscape utilizing the same method developed for ICM-Web. The default is 70% occupancy of the maximum number of allowed nucleosomes and a minimum linker length of 19 bps.

Shown in Figure 4, H. Varying the percent occupancy and minimum linker length determines how extended or condensed this non-uniform model will be.

#### Uniform

The Uniform option provides a uniform linker between all Masks. As in ICM-Web, the user can control the phase and linker length. The default value is a phase of 0 and linker length is 19. The first nucleosome begins at the very start of the DNA sequence. All successive nucleosomes are spaced 19 base pair from the previous one. This produces a chromatin fiber structure. As shown in the Figure 4, H, this uniform chromatin fiber is not necessarily straight even when the temperature is zero because the linker possesses sequence specific conformation and thermal properties.

#### Manual positioning

Users can manually control nucleosomes using the Nucleosome Widget. Clicking on one of the nucleosome “blocks” turns the block from transparent to green. The user can then drag the nucleosome to alter its position, change its type using a pull-down menu, or delete it. If there is sufficient room a nucleosome can be added to the left or right. Once the desired distribution of nucleosomes is achieved in the Nucleosome Energy Landscape, selecting “Position” from the drop down menu of build options will make the prescribed model and update the structure tracks to the genome browser.

#### Positioning from track

Finally, any single informatics track can be used to position nucleosomes. The chosen track may be experimentally or theoretically determined nucleosome positions or any combination of data uploaded by the user as track. To achieve this method of chromatin folding, the user clicks the desired track name and selects “Use position from track” in the “Position” drop down menu. Since nucleosome positions are displayed as a structure-track for any model created in G-Dash this track can be exported from the genome browser and uploaded later to restore a previous model with this modeling option. This technique can also be used to overlay a desired chromatin folding motif onto any segment of DNA.

#### Structure Relaxation

Steric clashes or knotting may occur for any model assembled in the material reference frame. For this purpose a minimization function is provided. When these problems occur, as indicated by red nucleosomes, users should click the “Minimize” button, to relax the model. If the problems can be resolved with a short minimization the user may assume the indicated 3D conformation can be physically realized. If the problems are not solved by the minimizer the model is likely not physically realizable.

#### Structural-Informatics

For the purposes of analyzing structure-function relationships it is beneficial to map informatics data onto a 3D model. G-Dash provides two methods of doing this. Individual nucleosomes can be selected in the Nucleosome Widget and assigned a specific color or an informatics track can be utilized to color all the DNA in a model. Selecting a track and using the TrackColor button in the control panel pushes informatics data onto the 3D model. JSmol’s command console can also be used to modify colors as desired, but it does not currently have direct access to the informatics data in the genome browser.

### From 3D Model to Informatics

The other usage modality of G-Dash is to extract informatics from a 3D structure and formats it as track data. This mode can be used to align an externally developed model with informatics data for the purpose of interpreting or validating the physical model or employed in a workflow for developing knowledge based potentials.

#### Structure Tracks

Whenever a model is made in G-Dash structural informatics tracks are generated and automatically updated, Figure 4, A. These tracks are highlighted in green in G-Dash and include helical parameter data, such as “Roll”, “Slide”, “Twist”, energy from the nucleosome energy landscape, and nucleosome occupancy data. These tracks all appear under the Modeling Data tab in Biodaliance’s track management tools.

#### Structure Upload

In the development version of G-Dash an externally developed 3D structure can be associated with a genome assembly using the upload button at the bottom of Control Panel. Currently only DiscoTech[65] based pdb models are allowed since the model has to be converted to an ICM compatible model. Any model for which 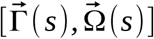 data can be computed should be supported. If the model explicitly contains 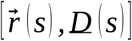 information or if the director frames can be systematically extracted from the model then the algorithms described in Methods can be used to compute structure tracks. If, as is the case with the DiscoTech based models, only 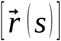 data is available, director frames must be determined from a TNB description. The DiscoTech based models pose the additional challenge that there are nine base pairs per bead. Uploading DiscoTech based models thus demonstrates an important proof of concept. TNB approach can be used to generate missing data. If the model contains sequence information it can be automatically aligned with a genome assembly. For the DiscoTech based models, which contain no sequence, the user chooses a starting location with the sequence selector (yellow bar) to associate the model with chromosome coordinates. This usage modality provides a powerful tool for interpreting modeling based studies.

## APPLICATION

As a sample application, G-Dash is utilized to studied the CHA1 promoter located on chromosome III of *saccharomyces cerevisiae (*sacCer3). This system is well studied and was used, along with the HIS3 and PHO5 promoters, to validate one of the first genome wide assays of nucleosome positions[66]. For this example experimentally determined and theoretically predicted nucleosome positions are used to make coarse-grained models of chromatin. We also consider the effects of the phase term in a uniform chromatin model.

### Generating User-defined Tracks

Nucleosome positioning information from [57] and hydroxyl radical cleavage data from [67] were converted to bigwig data format using the wigToBigWig converter provided by [58]. These tracks are uploaded as custom tracks in Biodalliance using its track management features by selecting the “Binary” option under the “Choose Files” button, and choosing the desired file. Clicking “Add” displays the bigwig data as a new track in G-dash(see supplemental video). Data from [57] is loaded by default for the sacCer3 assembly in the stable version of G-Dash. Data from [67] is loaded by default in the development version of G-Dash.

### Experimentally Determined Positions

Using chrIII: 15798..16880 as the CHA1 promoter region of interest and the experimentally determined nucleosome positions produces the model shown in Figure 4, G. Specifically, the track labeled “Occ” in G-Dash’s sacCer3 assembly has been used for the nucleosome positions. The resulting structure has both steric clash and knotting problems all of which are solved by using G-Dash’s “Minimize” feature, Figure 4, G. Similar to the results for the MMTV promoter computed by ICM-Web[45] it is clear that several irregularly spaced nucleosomes are insufficient to identify a chromatin fiber. Moreover, the 3D structure of the promoter is much more accessible than one may be lead to believe by spacing of nucleosomes as informatics, Figure 4, G. Assigning different conformations from the nucleosome superfamily to the individual nucleosomes or using chromatasome complexes rather than octasomes will certainly change conformation of the model; however, such changes will not produce a highly compact structure. This is true for any choice of nucleosome positions available from [57]. The PHO5 and HIS3 promoters yield similar results.

### Uniform

To achieve a highly compact structure one can ignore the nucleosome positioning data and uniformly space the nucleosomes, as is done for chrIII: 12871…16880 of sacCer3 in Figure 4, H. The resulting model has the general properties of a 30nm fiber. However when all nucleosomes are repositioned by 5 base pairs using G-Dash’s phase option the chromatin fiber bends, Figure 4, H. A similar phase dependent bending of uniform 30nm fiber models was obtained for the MMTV promoter using ICM-Web[45]. These models clearly demonstrate how the sequence dependent material properties (conformation and flexibility) of linker DNA can effect the structure of chromatin on longer length scales. In Figure 5, multiple loops occur in a uniform linker model of chromosome III and are observed to be a common feature of long length scale models produced by ICM-Web and G-Dash. The looping arises only from the sequence dependent material properties of DNA (conformation and flexibility).

**Figure 5:**
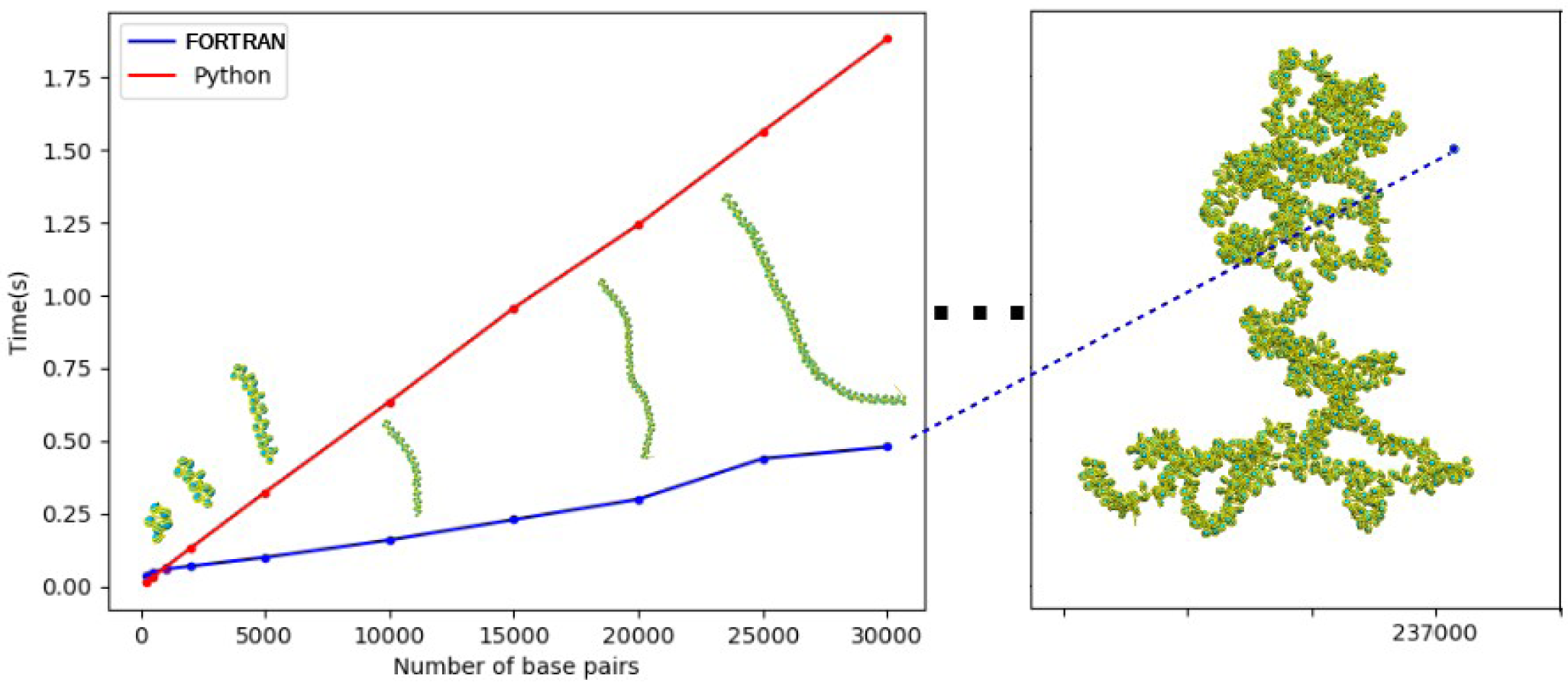
Timing Summary. Timing is reported for Python and FOTRAN versions of the compute kernel as a function of DNA length in base pairs. Both compute kernels exhibit linear costs as expected and can fold arbitrarily long sequences of DNA into extended or condensed models of DNA and chromatin. Reported time does not include transfering the model to the molecular viewer and rendering it. That timing depends strongly on network speed and congestion.

### G-Dash Scalability

The conversion between material frame and laboratory representations is fast. Compute time to fold 30,000bp is less than 0.5s. Folding all of chromosome III (over 237,000bp) requires less than 1.5s in the FORTRAN compute engine, Figure 5. Data for a Python based version is provided for comparison. For G-Dash the bottleneck is transferring and loading the molecule for display in JSmol. Speed up in G-Dash can be achieved by not transferring all the base pair data. In fact transferring only one bead per nucleosome rather than a bead for each base pair and the core to represent a nucleosome will yield an over 100 fold savings per nucleosome. These and other optimizations will allow megabase segments and larger to be managed interactively in a web format and small to medium chromosomes to be managed in computational workflows on a desktop workstation.

## DISCUSSION

Here, G-Dash has utilized experimentally determined and theoretically predicted nucleosome positions to make models of chromatin folding. G-Dash provides an approximate model. The structural details of both the positioned nucleosome and the uniform positioning models of CHA1 are likely inaccurate, but the gross scale features and interpretation are likely correct. 3D structures determined from nucleosome positioning data are much more extended and irregular than is typically appreciated. The overall conformation of chromatin can be radically changed by subtle changes in nucleosome positions, unless linker DNA deforms to preserve the spatial arrangement of the nucleosomes. Likewise, large changes in nucleosome positioning may yeild similar 3D conformations. Thus conserved patterns in 3D may exist that cannot be observed by consideration of nucleosome positions alone. A model of chromatin structure emerges in which nucleosomes serve dual functions: compacting the genome and regulating structural organization (chromatin looping) by masking and unmasking the material properties on DNA. These structure functions relationship can only be explored by unity informatics and 3D structural data.

Nucleosome positions and DNA materials properties alone are an incomplete model of chromatin. The obvious next step is inclusion of Hi-C, Micro-C and other cross-linking data as distance constraints in the 3D structures. This is a form of a knowledge-based potential that can be immediately realized by unifying structure and informatics in a genome dashboard. G-Dash is only a prototype of the genome dashboard concept that unifies structure and informatics approaches. We will continue to add features to G-Dash, but expect other genome dashboards to be developed. Using the design principals described here. There is nothing in the genome dashboard concept that limits its application to eukaryotes or chromatin folding.

## AKNOWLEDGEMENT

This effort was supported by the National Institute of General Medical Sciences of the National Institutes of Health under grant number 5 P20 GM103424-15 and 3 P20 GM103424-15S1. Partial support also received from NSF through cooperative agreement OIA-1541079 and the Louisiana Board of Regents.

## References

[1] Perisic, O. and Schlick, T. (2016). Computational strategies to address chromatin structure problems., Physical biology 13 : 035006.

[2] Fussner, E.; Ching, R. W. and Bazett-Jones, D. P. (2011). Living without 30nm chromatin fibers., Trends in biochemical sciences 36 : 1–6.

[3] Robinson, P. J. J.; An, W. and et al. (2008). 30 nm chromatin fibre decompaction requires both H4-K16 acetylation and linker histone eviction., Journal of molecular biology 381 : 816–825.

[4] Routh, A.; Sandin, S. and Rhodes, D. (2008). Nucleosome repeat length and linker histone stoichiometry determine chromatin fiber structure., Proceedings of the National Academy of Sciences of the United States of America 105 : 8872–8877.

[5] Davey, G. E.; Wu, B. and et al. (2010). DNA stretching in the nucleosome facilitates alkylation by an intercalating antitumour agent., Nucleic acids research 38 : 2081–2088.

[6] Schlick, T.; Hayes, J. and Grigoryev, S. (2012). Toward convergence of experimental studies and theoretical modeling of the chromatin fiber., The Journal of biological chemistry 287 : 5183–5191.

[7] Ozer, G.; Luque, A. and Schlick, T. (2015). The chromatin fiber: multiscale problems and approaches., Current opinion in structural biology 31 : 124–39.

[8] Bednar, J.; Garcia-Saez, I. and et al. (2017). Structure and Dynamics of a 197 bp Nucleosome in Complex with Linker Histone H1., Molecular cell 66 : 384-397.e8.

[9] Nikitina, T.; Norouzi, D. and et al. (2017). DNA topology in chromatin is defined by nucleosome spacing., Science advances 3 : e1700957.

[10] Belton, J.-M.; McCord, R. P. and et al. (2012). Hi-C: a comprehensive technique to capture the conformation of genomes., Methods (San Diego, Calif.) 58 : 268–276.

[11] Rao, S. S. P.; Huntley, M. H. and et al. (2014). A 3D map of the human genome at kilobase resolution reveals principles of chromatin looping., Cell 159 : 1665–1680.

[12] Mozziconacci, J. and Koszul, R. (2015). Filling the gap: Micro-C accesses the nucleosomal fiber at 100-1000 bp resolution., Genome biology 16 : 169.

[13] Rotem, A.; Ram, O. and et al. (2015). Single-cell ChIP-seq reveals cell subpopulations defined by chromatin state., Nature biotechnology 33 : 1165–1172.

[14] Hsieh, T.-H. S.; Fudenberg, G. and et al. (2016). Micro-C XL: assaying chromosome conformation from the nucleosome to the entire genome., Nature methods 13 : 1009–1011.

[15] Belaghzal, H.; Dekker, J. and Gibcus, J. H. (2017). Hi-C 2.0: An optimized Hi-C procedure for high-resolution genome-wide mapping of chromosome conformation., Methods (San Diego, Calif.) 123 : 56–65.

[16] Marti-Renom, M. A.; Almouzni, G. and et al. (2018). Challenges and guidelines toward 4D nucleome data and model standards., Nature genetics 50 : 1352–1358.

[17] Chang, P.; Gohain, M. and et al. (2018). Computational Methods for Assessing Chromatin Hierarchy., Computational and structural biotechnology journal 16 : 43–53.

[18] Auton, A.; Brooks, L. D. and et al. (2015). A global reference for human genetic variation., Nature 526 : 68–74.

[19] Sudmant, P. H.; Rausch, T. and et al. (2015). An integrated map of structural variation in 2,504 human genomes., Nature 526 : 75–81.

[20] Consortium, E. P. (2012). An integrated encyclopedia of DNA elements in the human genome., Nature 489 : 57–74.

[21] Pazin, M. J. (2015). Using the ENCODE Resource for Functional Annotation of Genetic Variants., Cold Spring Harbor protocols 2015 : 522–536.

[22] Diehl, A. G. and Boyle, A. P. (2016). Deciphering ENCODE., Trends in genetics : TIG 32 : 238–249.

[23] Dekker, J.; Belmont, A. S. and et al. (2017). The 4D nucleome project., Nature 549 : 219–226.

[24] Cowper-Sal lari, R.; Zhang, X. and et al. (2012). Breast cancer risk-associated SNPs modulate the affinity of chromatin for FOXA1 and alter gene expression., Nature genetics 44 : 1191–8.

[25] Lott, P. C. and Carvajal-Carmona, L. G. (2018). Resolving gastric cancer aetiology: an update in genetic predisposition., The lancet. Gastroenterology & hepatology 3 : 874–883.

[26] Xiong, J. and Zhao, W.-L. (2018). Advances in multiple omics of natural-killer/T cell lymphoma., Journal of hematology & oncology 11 : 134.

[27] Barbara, M.; Scott, A. and Alkhouri, N. (2018). New insights into genetic predisposition and novel therapeutic targets for nonalcoholic fatty liver disease., Hepatobiliary surgery and nutrition 7 : 372–381.

[28] Perkel, J. M. (2017). Plot a course through the genome., Nature 549 : 117–118.

[29] Pasi, M.; Maddocks, J. H. and et al. (2014). muABC: a systematic microsecond molecular dynamics study of tetranucleotide sequence effects in B-DNA., Nucleic acids research 42 : 12272–83.

[30] Korolev, N.; Lyubartsev, A. P. and Nordenskiöld, L. (2018). A systematic analysis of nucleosome core particle and nucleosome-nucleosome stacking structure., Scientific reports 8 : 1543.

[31] Simo, J. C. and Vu-Quoc, L. (1991). A geometrically-exact rod model incorporating shear and torsion-warping deformation, International Journal of Solids and Structures 27 : 371–393.

[32] Struik, D. J., (1988). Lectures on Classical Differential Geometry: Second Edition. Dover Publications,.

[33] Calladine, C. R. and Drew, H. R. (1986). Principles of sequence-dependent flexure of DNA., Journal of molecular biology 192 : 907–918.

[34] Mergell, B.; Ejtehadi, M. R. and Everaers, R. (2003). Modeling DNA structure, elasticity, and deformations at the base-pair level., Physical review. E, Statistical, nonlinear, and soft matter physics 68 : 021911.

[35] Becker, N. B. and Everaers, R. (2007). From rigid base pairs to semiflexible polymers: coarse-graining DNA., Physical review. E, Statistical, nonlinear, and soft matter physics 76 : 021923.

[36] Fathizadeh, A.; Eslami-Mossallam, B. and Ejtehadi, M. R. (2012). Definition of the persistence length in the coarse-grained models of DNA elasticity., Physical review. E, Statistical, nonlinear, and soft matter physics 86 : 051907.

[37] Gonzalez, O.; Petkevičiūtė, D. and Maddocks, J. H. (2013). A sequence-dependent rigid-base model of DNA., The Journal of chemical physics 138 : 055102.

[38] Dickerson, R. (1989). Definitions and nomenclature of nucleic acid structure parameters, J Biomol Struct Dyn 6 : 627–634.

[39] Dickerson, R. (1989). Definitions and nomenclature of nucleic acid structure component, Nucleic Acids Res 17 : 1797–1803.

[40] el Hassan, M. A. and Calladine, C. R. (1995). The assessment of the geometry of dinucleotide steps in double-helical DNA; a new local calculation scheme., Journal of molecular biology 251 : 648– 64.

[41] Petkeviciute, D.; Pasi, M.; Gonzalez, O. and Maddocks, J. H. (2014). cgDNA: a software package for the prediction of sequence-dependent coarse-grain free energies of B-form DNA., Nucleic acids research 42 : e153.

[42] Olson, W. K.; Gorin, A. A. and et al. (1998). DNA sequence-dependent deformability deduced from protein-DNA crystal complexes., Proc Natl Acad Sci U S A 95 : 11163–11168.

[43] Czapla, L.; Swigon, D. and Olson, W. K. (2006). Sequence-Dependent Effects in the Cyclization of Short DNA., Journal of chemical theory and computation 2 : 685–695.

[44] Lavery, R.; Zakrzewska, K. and et al. (2010). A systematic molecular dynamics study of nearest-neighbor effects on base pair and base pair step conformations and fluctuations in B-DNA., Nucleic Acids Res 38 : 299–313.

[45] Stolz, R. C. and Bishop, T. C. (2010). ICM Web: the interactive chromatin modeling web server., Nucleic acids research 38 : W254–61.

[46] Lu, X.-J. and Olson, W. K. (2003). 3DNA: a software package for the analysis, rebuilding and visualization of three-dimensional nucleic acid structures., Nucleic Acids Res 31 : 5108–5121.

[47] Lavery, R.; Moakher, M. and et al (2009). Conformational analysis of nucleic acids revisited: Curves+., Nucleic Acids Res 37 : 5917–5929.

[48] Babcock, M. S.; Pednault, E. P. and Olson, W. K. (1993). Nucleic acid structure analysis: a users guide to a collection of new analysis programs., Journal of biomolecular structure & dynamics 11 : 597–628.

[49] D.A. Case, I. B.-S. and et al. (2018). AMBER 2018, University of California, San Francisco..

[50] Babcock, M. S.; Pednault, E. P. and Olson, W. K. (1994). Nucleic acid structure analysis. Mathematics for local Cartesian and helical structure parameters that are truly comparable between structures., Journal of molecular biology 237 : 125–156.

[51] Olson, W. K.; Bansal, M. and et al. (2001). A standard reference frame for the description of nucleic acid base-pair geometry., Journal of molecular biology 313 : 229–37.

[52] Korolev, N.; Lyubartsev, A. P. and Nordenskiold, L. (2010). Cation-induced polyelectrolyte-polyelectrolyte attraction in solutions of DNA and nucleosome core particles., Advances in colloid and interface science 158 : 32–47.

[53] Korolev, N.; Zhao, Y. and et al. (2012). The effect of salt on oligocation-induced chromatin condensation., Biochemical and biophysical research communications 418 : 205–10.

[54] Dijkstra, E. W. (1974). Programming as a discipline of mathematical nature, Am. Math. Monthly 81 : 608–612.

[55] Down Thomas A. M. P. and Hubbard., T. J. P. (2011). Dalliance: Interactive Genome Viewing on the Web., Bioinformatics 27.6 : 889–890.

[56] Biodalliance Getting Started. http://www.biodalliance.org/started.html,.

[57] Jiang, C. and Pugh, B. F. (2009). A compiled and systematic reference map of nucleosome positions across the Saccharomyces cerevisiae genome., Genome biology 10 : R109.

[58] Kent WJ Z. A. and et al. (2010 Sep 1). BigWig and BigBed: enabling browsing of large distributed data sets., Bioinformatics. 26(17) : 2204–7.

[59] Bishop, T. C. (2008). Geometry of the nucleosomal DNA superhelix., Biophysical journal 95 : 1007–17.

[60] Zlatanova, J.; Bishop, T. C. and et al. (2009). The nucleosome family: dynamic and growing., Structure (London, England : 1993) 17 : 160–71.

[61] Davey, C. A.; Sargent, D. F. and et al. (2002). Solvent mediated interactions in the structure of the nucleosome core particle at 1.9 a resolution., Journal of molecular biology 319 : 1097–1113.

[62] Smith, J. A.; Romanus, M.; et al. (2013). Scalable Online Comparative Genomics of Mononucleosomes: A BigJob, : 23:1-23:8.

[63] Thibault, J. C.; Cheatham, T. E. and Facelli, J. C. (2014). iBIOMES Lite: summarizing biomolecular simulation data in limited settings., J Chem Inf Model 54 : 1810–1819.

[64] xTMB-IBIOMES. http://dna.engr.latech.edu/∼ibiomes/.

[65] Zhang, Q.; Beard, D. A. and Schlick, T. (2003). Constructing irregular surfaces to enclose macromolecular complexes for mesoscale modeling using the discrete surface charge optimization (DISCO) algorithm., Journal of computational chemistry 24 : 2063–2074.

[66] Field, Y.; Kaplan, N. and et al. (2008). Distinct modes of regulation by chromatin encoded through nucleosome positioning signals., PLoS computational biology 4 : e1000216.

[67] Chiu, T.-P.; Yang, L. and et al. (2015). GBshape: a genome browser database for DNA shape annotations., Nucleic acids research 43 : D103–D109.

